# Life history of Dactylopius opuntiae (Hemiptera: Dactylopiidae) on Moroccan resistant cactus germplasm

**DOI:** 10.1101/2021.07.23.453565

**Authors:** Mohamed El Aalaoui, Mohamed Sbaghi

## Abstract

The important damages caused by *Dactylopius opuntiae* (Hemiptera: Dactylopiidae) to cactus crops around the world require an integrated pest management (IPM) approach, based on the combination of several techniques (varietal resistance, biological, chemical methods, etc). In this sense, this study evaluated the resistance of 10 Moroccan cactus genotypes to *D. opuntiae* in order to characterize the expression of antixenosis and/or antibiosis. Antixenosis was accessed in the greenhouse and in the laboratory (26± 2°C) using choice and non-choice tests with 1^st^ instar nymphs. Aakria and Cherratia showed a strong antixenosis effect towards *D. opuntiae* (0-0.3 *D. opuntiae* alive 30 after infestation). For antibiosis assessment, 30 1^st^ instar nymphs were confined on cladodes of the 10 selected genotypes under the same laboratory conditions to allow their development, as well as the life cycle performance and behavior of *D. opuntiae* on the 10 selected cactus genotypes, were evaluated under greenhouse conditions. No influence of genotypes on insect oviposition was observed, indicating that the mealybug does not prefer any genotypes over the others for oviposition. The mealybug failed to develop on genotypes Aakria and Cherratia and did not grow beyond the young female stage on all other resistant genotypes tested. Similarly, first instar nymphs fed on genotypes Marjana, Melk Zhar, and A200 died without reaching the second instar nymph stage. In addition, all genotypes tested had a negative effect on nymph viability (<24%), indicating resistance (antibiosis and/or antixenosis) to the cactus scale. These cactus genotypes may all be useful in breeding programs focused on cactus resistance to mealybugs.

## Introduction

Worldwide, approximately 130 genera and 2,000 species belonging to the family Cactaceae (order Caryophyllales) have been recorded, almost all of which are native to the Americas. Mexico is where the center of diversity for cacti in the world is based [1, 2]. Cacti occur in a wide variety of sizes and shapes, from the smallest species measuring 9 mm in diameter to the largest measuring nearly 20 m [3]. Cactaceae generally have a diploid chromosome system of 2n = 22, but many species in the family Opuntioideae are known to be polyploid [3].

The cactus is well known as a multipurpose crop with an important ecological and economic role. Indeed, cacti can be used as fodder, medicinal plants, for human consumption (vegetables and fruits), and as ornamental plants [4]. It can also be used as a food additive, nutritional supplement, and for cosmetic and pharmaceutical purposes [4]. The cactus is grown commercially as a fruit plant in only five countries: Chile, Mexico, Italy, the United States, and South Africa [5]. Cactus fruits are very nutritious. They are rich in vitamins, minerals, proteins, phenolic compounds, and other elements with high nutritional value [6, 7]. This crop is known as a good indicator of harmfulness (Nobel, 1994) and as a living fence to protect agricultural areas [8]. However, this crop is subject to a number of biotic constraints such as fungal diseases, lepidopterans, gastropods, cactus mealybugs (*Diaspis echinocacti* and *Dactylopius opuntiae*) that have very dangerous impacts on the yield and the sustainability of cactus cultivation even around the world. Indeed, *D. opuntiae* (Cockerell) is one of the eleven species belonging to the monogenic family Dactylopiidae [9], considered among the main pests of *Opuntia ficus-indica* (L.) Miller (Caryophyllales: Cactaceae) and other cultivated and wild Opuntia species in many countries in the world [10, 11, 12]. The mealybug (*D. opuntiae*) has also been known as a biological agent for the control of Opuntia in countries where they behave as invasive plants [13]. *Dactylopius opuntiae* is a sap-sucking insect that can have a strong negative impact on both prickly pear production for fresh consumption and cladodes used as fodder for livestock [14]. The mealybug tends to form colonies of variable size on cladodes, which in some cases are completely covered by the insect [15, 16], which triggers the desiccation and loss of cladodes, premature drop of fruits and total death of the cactus plant [17]. Thus, the severe damage caused by this insect requires an integrated pest management (IPM) approach, based on the combination of several techniques, including genetic, mechanical, physical, biological, chemical, etc. methods [18, 19, 20], in order to obtain better results in the control of this pest.

For Morocco, the prickly pear has been introduced since the 16th century [21]. The cactus is very common in most regions of the country where the crop has become important. But unfortunately with the appearance for the first time in Morocco of the wild mealybug of cactus “*Dactylopius opuntiae*” in 2014, the cactus sector found itself in front of a very big scourge. The rapid and unpredictable spread of this mealybug from the first outbreak to other cactus production areas of the country has led to the destruction of several hedgerows and cactus plantations where the mealybug has devastated thousands of hectares and kilometers of cactus plantations, causing huge socio-economic and environmental losses. Similar cases were reported by Lopes et al. in 2009 where *D. opuntiae* attacks on a cactus forage species, *Opuntiae ficus indica*, in Brazil, resulted in the loss of 100,000 ha, valued at 25 million dollars [22].

Considering the urgency of the mealybug, and to avoid the spread of this epidemic, the Ministry of Agriculture, Maritime Fishing, Rural Development, and Water and Forests-Morocco, put in place a major emergency plan for the control of this mealybug in 2016. This plan also included a research program covering the most important management elements such as host plant resistance [23], pesticides, beneficial insects [24, 25, 26, 27, 28], and biopesticides [29]. Of all the investigated research pathways, the identification of ten mealybug-resistant genotypes is the part that offers great hope for the revival of the cactus industry at the present time [23]. Resistant genotypes are showing positive results in Brazil as well. Matos et al. (2021) reported that the best alternative for the cultivation of cactus in regions attacked by this insect is to plant cultivars resistant to the carmine scale [30]. Thus, to provide effectively and simultaneously environmentally and human-friendly alternatives, the use of resistant genotypes can be a valuable strategy for integrated pest management (IPM) [31].

Identifying and characterizing the defense mechanisms of the resistant cactus would be useful to researchers, as it would allow the development of molecular markers that could be used for targeted selection. However, resistance can be a complex phenomenon. Painter (1951) describes three categories of resistance: tolerance, non-preference (or antixenosis), and antibiosis [32]. In plant tolerance, the insect attacks the plant, but the plant has the ability to recover from the wound, and the insect’s biological performance is not altered and its behavior is not negatively affected [33]. Antixenosis describes the category of plant defense where plants have chemical or physical characteristics that make it less probable that an herbivore will use this plant as a host [32, 34]. Insects use olfactory, gustatory, tactile, and visual cues to make host selection decisions, and anti-xenotic properties have been described in numerous studies of insect-plant systems for all of these cues, most of which have been described in agricultural systems [31]. Antibiosis can be defined as the category of plant resistance in which plants employ mechanisms that deleteriously affect herbivores once they have chosen to feed on this plant [32, 35]. Plants use a variety of antibiosis mechanisms, including toxic secondary metabolites such as chemicals, proteins, mechanical defenses, or combinations of these [36]. The antibiotic effects of these mechanisms can range from mild to lethal and even if an individual survives, they may suffer crippling effects such as reduction in body size, fecundity, and prolonged developmental periods [31]. In addition, induced responses are critical components of antibiosis, where certain cues such as herbivore feeding, salivary enzymes, and plant hormones cause the expression of certain defenses [31, 37].

Considering the enormous damage caused by this very devastating pest to cacti in Morocco and worldwide and considering that this plague continues its progression in different cactus production areas of the country and also the need to identify more sustainable control methods, the present study aimed to characterize the resistance of the 10 cactus genotypes identified as resistant to *D. opuntiae* in Morocco using antexenosis and antibiosis tests in laboratory and greenhouse.

## Material and Methods

### Site of study and plant material

The study was conducted at the Agricultural Technical Institute of Khmiss Zemamra-Doukkala (2020-2021). The Doukkala region extends between latitudes 32°15 and 33°15 North and longitudes 7°55 and 9°15 West. It straddles the provinces of El Jadida and Safi. It is limited to the northeast by the Chaouia, southwest by the region of Abda, west by the Atlantic Ocean, and southeast by the massifs of R’hamna. The locality of Zemamra is located in a semi-arid ecological zone where annual rainfall varies between 112.6 mm and 607 mm. The annual average of 30 years is 330 mm. The temperature varies from -1 °C to 45 °C.

The ten cactus genotypes tested in this study were brought collected from an INRA (National Institute for Agricultural Research) national cactus collection. Eight of these resistant genotypes (Karama, Ghalia, Belara, Marjana, Melk Zhar, Cherratia, Angad, and Aakria) have been identified as resistant to *D. opuntiae* and are already registered in the Catalogue Officiel of Cactus in Morocco [23] (Table 1).

**Table 1.** List of cactus genotypes resistant to *D. opuntiae* and registered in the catalog officiel of cactus in Morocco [23]

| Genotypes | Origin | Characteristics of cladodes | Fruit characteristics |
| --- | --- | --- | --- |
| Marjana | Dchira- Inezgane - Morocco | Intermec cladodes (without spines) for use as fodder | Fruit with light purple flesh, juicy and very sweet |
| Belara | Dchira - Inezgane - Morocco | Intermec cladodes (without thorns) with good forage quality | Fruit with white flesh, juicy and very sweet |
| Karama | Dchira - Inezgane - Morocco | Spiny cladodes with very good forage quality | Fruit with red flesh, very sweet and tasty |
| Ghalia | Dchira - Inezgane - Morocco | Thorny cladodes of good quality and very rich in nitrogen for livestock | Fruit with very good organoleptic quality, rich in vitamins and antioxidants, low in acidity, and very sweet |
| Angad | Oujda - Morocco | Very thorny cladodes with good quality and very rich in nitrogen for livestock | Fruit with dark-purple flesh, sweet, tasty, and very rich in vitamin C and antioxidants |
| Aakria | Bouknadel - Morocco | Thorny cladodes with good forage quality | Red fruit of small size, too acidulous, not very sweet, and appreciated in the off-season by diabetics mainly |
| Melk Zhar | Irradiation *O. robusta* - Morocco | Very thorny cladodes with good quality and very high nitrogen content for livestock | Fruit with very good organoleptic quality, rich in vitamins and antioxidants, low in acidity |
| Cherratia | Bouznika-Morocco | Very thorny cladodes with good quality and very rich in nitrogen for livestock | Fruit with very good organoleptic quality, low acidity, rich in vitamins and antioxidants, and very sweet |

### The cactus mealybug colony

A strain colony of *D. opuntiae* was started with infested cladodes of *Opuntia ficus-indica* (L.) Miller, 1768) collected from fields in the locality of Zemamra (32°37’48” N, 8°42’0” W) in the Casablanca-Settat region, Morocco and maintained under laboratory conditions (26± 2°C, 60± 10% RH, and 12h photophase), using a modified version of the method described by Aldama-Aguilera and Llanderal-Cazares (2003) [38]. Infested cladodes were placed in entomological cages (80***×*** 80***×***80 cm) consisting of a wooden frame covered with mesh fabric to allow ventilation. Each cladode was punctured at the basal end by a wooden stake, left to heal for 24 hours under laboratory conditions (26± 2°C, 60± 10% RH), and then suspended vertically from metal grids; other uninfested cladodes collected from the same site (Zemamra) were placed horizontally beneath for nymphs that had detached on the vertical cladodes. Healthy, uninfested cladodes were introduced weekly into the cages to maintain the colony. Portions of cotton moistened with distilled water were placed in the bottoms of inverted Petri dishes (14.5 cm diameter) and introduced into the cages to maintain humidity. In order to increase the insect numbers and monitor its age, the first instar nymphs of *D. opuntiae* (24 hours old) were transferred to another cage with the same characteristics as described above to complete their development.

### Antixenosis test

#### Under laboratory

Under laboratory conditions at 26± 2°C, 60± 10% RH, non-preference (antixenosis) assays were evaluated in free choice and non-choice test studies against the first nymphal stages of the scale insect. To investigate the possibility of non-preference interaction in choice tests, the ten genotypes tested cladodes (one-year-old) (n = 3) with a susceptible control were placed and arranged in a circle in entomological cages (80×80×80 cm) with the same characteristics as described above (Cactus mealybug colony section). Five cladodes of *Opuntia ficus-indica* (L.) heavily infested with *D*.*opuntiae* were placed in the center of each cage and equidistant from the cladodes. The number of alive insects (attracted) in each cladode was recorded 1, 3, 15, 30 days after infestation using a binocular loupe (Motic). This test had 20 replicates in a completely randomized design. For the no-choice tests, we followed the same procedures as for the choice tests but this time the cladodes were placed separately according to genotype in entomological cages (80× 80×80cm). Study design and replication were the same for the free-choice test. The number of insects alive was measured at 1, 3, 15, and 30 days after infestation, similar to what was described for the free choice test.

#### Under greenhouse

To assess the preference of *D. opuntiae* among 10 cactus genotypes, we performed a multiple-choice test, using a modified version of the methodology used with *Aphys glycines* Matsumura (Hemiptera: Aphididae) [39, 40] and *Dichelops melacanthus* Dallas (Hemiptera: Pentatomidae) [41]. The resistant genotypes’ with susceptible control cladodes (one-year-old) were planted in normal polarity in a plastic pot (33 cm diameter by 12 cm height), filled with a mixture of fine sand (2/3) and peat (1/3), and grown until the plants reached the stage of three to five cladodes. Then the plants were arranged in completely randomized rows (1 m between rows, with 5 cm spacing between plants) under the greenhouse. Plants were irrigated as needed. Between the lines and approximately 50 cm from each pot, an *Opuntia ficus indica* (L.) cladode that was highly infested with first and second instar nymphs of *D. opuntiae* (1-15 days) was placed. These stages were chosen because the nymphs do not fly and the plots used had no cover on top. The ten genotypes were evaluated using a completely randomized design with twenty replicates. The number and stage of insects per plant were recorded at 1, 3, 15, and 30 days after infestation with help of a handloup. The semi-field daily temperature ranged 8-30 °C during this study and was recorded using thermograms, based on 6 measurements made with intervals of 2 h. The night temperature was determined from the 3 lowest daily values.

### Antibiosis Test

#### Under laboratory

The test was performed with cladodes of similar age and under the same laboratory conditions used in the previous test. However, in this study, cladodes of different genotypes were placed individually inside entomological cages (80× 80×80cm) (similar to those described previously) and each cladode was infested with 2 mature females of *D*.*opuntiae* (egg production stage). All cladodes were infested with mature females of similar age and weight. After infestation, the total number of eggs laid on each cladode was recorded. Eggs were placed in labeled Petri dishes (14.5 cm diameter) and observed daily and the date of hatching was recorded to determine the incubation period. Hatchability (%) was calculated using the equation from Abbas et al. (2012): Hatchability = All neonates/All eggs [42]. After hatching, 30 first instar nymphs were left on each cladode and allowed to develop. First instar nymph viability (i.e., successful development to second instar nymph) and duration of each stage reached was recorded. The morphology of the different life cycle stages developed on each genotype and the behavior of the insects were examined using a binocular loupe (Motic) (results no-showed in this manuscript). The mealybug stages reached on each genotype weight (mg) was measured using an electronic balance with a precision of 0.001 mg (OHAUS CORPORATION, USA). In this experiment, each cladode represented a replicate, with 20 replicates per genotype, in a completely randomized design.

#### Under greenhouse

The life cycle performance and behavior of *D. opuntiae* on 10 selected cactus genotypes were evaluated in a greenhouse under the same temperature conditions as for the Antixenosis test (8-30 °C). The ten genotypes with a susceptible control were evaluated in a completely randomized design with 20 replicates. Cactus plants offered to the mealybug were in the 3-5 cladodes stage.

The genotypes tested cladodes (one-year-old) were planted in plastic pots (similar in volume to those used in the antixenosis test), filled with the same substrate as described above. Cactus plants were infested with *Opuntiae ficus indica* (L.) cladodes heavily infested with *D. opuntiae* 1st instar nymphs and placed close to the axis of each plant to allow for movement and attachment of nymphs to appropriate zones on the plant. The poles were evaluated daily to assess the duration of each stage reached.

#### Morphological characterization of plants

Spine and cladode measurements are important informative characters for the taxonomy of Opuntia species [43]. Therefore, the methodological parameters used in this study were adapted from similar methods in previous work on Opuntia morphology by Mosco (2009), Peharec et al. (2010), and Musengi et al. (2021) [43, 44, 45]. The parameters measured were: total number of spines per cladode, cladode surface area, number of areoles per cladode, and number of areoles per plant.

#### Statistical analyses

The normality of the Antixenosis and Antibiosis assays data were evaluated using the Shapiro-Wilk W test. One-way analysis of variance (ANOVA) was used for the analysis of the number of insects attracted to the different cactus genotypes, insect development, and nymphal viability trials to compare the development of *D. opuntiae* among the host plants under laboratory and greenhouse conditions with regard to the duration of instars and mortality rates. These data were examined using analysis of variance, and means were compared with Tukey’s LSD test (α = 0.05).

## Results

### Antixenosis test

#### Under laboratory

Significant differences were observed among cactus genotypes in the four periods of attractiveness assessment with *D*.*opuntiae* nymphs (Table 2). Generally for the two tests performed (free choice and non-choice), At 1, 3, and 15 days after infestation, the genotype Aakria was the least attractive. However, at 30 days after release, no differences were observed among the resistant genotypes tested, and the control genotype (405-424 nymphs) was the most infested.

**Table 2.** Mean (±SE) number of *Dactylopius opuntiae* on cactus genotypes in different periods in an antixenosis experiment under laboratory conditions (26± 2°C, 60± 10% RH, and 12h photophase)

| Genotypes | Number of attracted insects in free-choice test<sup>a</sup> |  |  |  | Number of attracted insects in no-choice test<sup>a</sup> |  |  |  |
| --- | --- | --- | --- | --- | --- | --- | --- | --- |
|  | 1 d | 3 d | 15 d | 30 d | 1 d | 3 d | 15 d | 30 d |
| Aakria | 5.9<br>$\pm 1.1$ e | 2.8<br>$\pm 1.1$ e | 2.3<br>$\pm 0.6$ c | 0.0<br>$\pm 0.0$ b | 7.0<br>$\pm 1.2$ f | 4.0<br>$\pm 1.1$ e | 2.8<br>$\pm 0.8$ d | 0.3<br>$\pm 0.4$ b |
| Cherratia | 15.7<br>$\pm 1.5$ d | 10.8<br>$\pm 1.5$ de | 9.6<br>$\pm 1.2$ bc | 0.2<br>$\pm 0.4$ b | 28.1<br>$\pm 3.8$ e | 21.4<br>$\pm 1.6$ d | 12.9<br>$\pm 2.9$ c | 0.3<br>$\pm 0.5$ b |
| Marjana | 16.0<br>$\pm 1.5$ d | 11.6<br>$\pm 1.9$ cde | 9.9<br>$\pm 1.3$ bc | 0.3<br>$\pm 0.5$ b | 35.3<br>$\pm 2.6$ de | 30.2<br>$\pm 2.8$ cd | 14.3<br>$\pm 2.4$ c | 0.4<br>$\pm 0.5$ b |
| Melk Zhar | 26.0<br>$\pm 1.9$ c | 15.8<br>$\pm 1.5$ bcd | 8.7<br>$\pm 1.2$ bc | 0.5<br>$\pm 0.6$ b | 44.1<br>$\pm 2.5$ cd | 38.5<br>$\pm 2.4$ bc | 15.3<br>$\pm 2.7$ c | 0.4<br>$\pm 0.6$ b |
| Angad | 26.9<br>$\pm 2.5$ c | 16.5<br>$\pm 1.6$ bcd | 9.5<br>$\pm 1.9$ bc | 0.7<br>$\pm 0.8$ b | 47.3<br>$\pm 2.4$ c | 40.9<br>$\pm 3.3$ bc | 16.8<br>$\pm 2.6$ c | 0.5<br>$\pm 0.6$ b |
| B176 | 30.4<br>$\pm 1.7$ c | 21.2<br>$\pm 1.8$ bcd | 10.9<br>$\pm 1.6$ b | 0.8<br>$\pm 0.8$ b | 51.8<br>$\pm 2.7$ bc | 41.1<br>$\pm 1.7$ bc | 18.9<br>$\pm 3.1$ bc | 0.7<br>$\pm 0.7$ b |
| B180 | 30.5<br>$\pm 2.0$ c | 22.2<br>$\pm 1.9$ bc | 11.2<br>$\pm 1.8$ b | 0.8<br>$\pm 0.8$ b | 51.2<br>$\pm 2.7$ bc | 41.7<br>$\pm 1.9$ bc | 19.8<br>$\pm 2.9$ bc | 0.8<br>$\pm 0.8$ b |
| Karama | 32.8<br>$\pm 1.6$ bc | 22.2<br>$\pm 2.4$ bc | 11.2<br>$\pm 1.5$ b | 0.9<br>$\pm 0.8$ b | 53.2<br>$\pm 1.7$ bc | 43.1<br>$\pm 1.8$ b | 20.4<br>$\pm 2.7$ bc | 0.9<br>$\pm 0.8$ b |
| Ghalia | 32.9<br>$\pm 1.5$ bc | 23.3<br>$\pm 2.1$ bc | 11.4<br>$\pm 2.0$ b | 1.1<br>$\pm 0.8$ b | 53.1<br>$\pm 1.5$ bc | 44.2<br>$\pm 2.3$ b | 20.7<br>$\pm 2.5$ bc | 1.2<br>$\pm 0.9$ b |
| Belara | 39.3<br>$\pm 2.7$ b | 25.5<br>$\pm 2.6$ b | 15.4<br>$\pm 2.3$ b | 1.3<br>$\pm 1.1$ b | 60.2<br>$\pm 2.5$ b | 46.6<br>$\pm 4.2$ b | 26.9<br>$\pm 3.8$ b | 1.5<br>$\pm 0.9$ b |
| Control | 176.3<br>$\pm 21.8$ a | 319.6<br>$\pm 35.2$ a | 361.7<br>$\pm 24.4$ a | 405.8<br>$\pm 19.9$ a | 197.0<br>$\pm 34.8$ a | 340.6<br>$\pm 38.3$ a | 374.0<br>$\pm 29.6$ a | 424.4<br>$\pm 32.9$ a |
| Statistics | $F =$<br>937.57<br>df = 10,<br>$P <$<br>0.0001 | $F =$<br>1448.45<br>df = 10,<br>$P <$<br>0.0001 | $F =$<br>3982.03<br>df = 10,<br>$P <$<br>0.0001 | $F = 8167.83$<br>df = 10,<br>$P < 0.0001$ | $F =$<br>410.79<br>df = 10,<br>$P <$<br>0.0001 | $F =$<br>1242.07<br>df = 10,<br>$P <$<br>0.0001 | $F =$<br>2688.69<br>df = 10,<br>$P <$<br>0.0001 | $F =$<br>3311.98<br>df = 10,<br>$P <$<br>0.0001 |

Means within a column followed by the same letters are not significantly different according to Tukey's LSD test at $\alpha = 0.05$

#### Under greenhouse

Regarding the preference of *D. opuntiae* nymphs on different cactus genotypes, at 24 h after release, Aakria (23.2 nymphs/plant) and Cherratia (45.1 nymphs/plant) genotypes were the least infected by *D. opuntiae* 1^st^ instars. No differences were observed among the different cactus resistant genotypes tested at 3, 15, and 30 days after the release of the scale pest nymphs. The susceptible control genotype was the most infested (Table 3).

**Table 3.** Mean (±SE) number of *Dactylopius opuntiae* alive on cactus genotypes in different periods in an antixenosis experiment under semi-field conditions (i.e., choice test experiment)

| Genotypes | Time<sup>a</sup> |  |  |  |  |  |  |  |
| --- | --- | --- | --- | --- | --- | --- | --- | --- |
|  | 1 d |  | 3 d |  | 15 d |  | 30 d |  |
| Aakria | 23.2±4.6 | f | 10.6±3.5 | b | 6.0±1.7 | b | 0.0±0.0 | b |
| Cherratia | 45.1±3.9 | ef | 30.9±4.6 | b | 11.5±2.5 | b | 0.3±0.5 | b |
| Marjana | 62.7±5.8 | de | 11.6±1.9 | b | 11.2±1.8 | b | 0.7±0.8 | b |
| Melk Zhar | 77.0±6.0 | cd | 30.5±3.4 | b | 11.5±1.9 | b | 0.8±0.9 | b |
| Angad | 79.0±6.4 | cd | 32.1±4.0 | b | 11.9±2.0 | b | 0.9±1.0 | b |
| B176 | 89.4±5.5 | bcd | 41.7±2.5 | b | 12.1±2.0 | b | 1.1±1.0 | b |
| B180 | 89.8±7.6 | bcd | 43.9±3.7 | b | 13.0±2.3 | b | 1.2±1.0 | b |
| Karama | 94.2±5.2 | bc | 41.7±6.0 | b | 13.5±2.0 | b | 1.4±1.0 | b |
| Ghalia | 95.5±4.5 | bc | 41.6±6.5 | b | 13.7±1.9 | b | 1.4±1.0 | b |
| Belara | 112.8±11.0 | b | 49.6±5.4 | b | 14.2±1.7 | b | 1.7±0.9 | b |
| Control | 702.2±85.9 | a | 1592.8±180.0 | a | 1804.5±120.9 | a | 2029.0±99.6 | a |
| Statistics | $F = 1022.25$ | | $F = 1492.85$ | | $F = 4382.44$ | | $F = 8283.89$ | |
| | df = 10, $P < 0.0001$ | | df = 10, $P < 0.0001$ | | df = 10, $P < 0.0001$ | | df = 10, $P < 0.0001$ | |

Means within a column followed by the same letters are not significantly different according to Tukey's LSD test at $\alpha = 0.05$

### Antibiosis Test

#### Under laboratory

No significant differences were observed among genotypes on the number of eggs per cladode, incubation period, and hatchability percentage (Table 4). The scale pest does not reach to develop definitively on the genotypes Aakria and Cherratia, and doesn’t get beyond the young female stage in all the other tested resistant genotypes. For the other genotypes, the development time of *D. opuntiae* nymphs was significantly affected by different cactus genotypes in 1^st^ and 2^nd^ instars.

**Table 4.** The developmental parameters of *Dactylopius opuntiae* on cactus genotypes in an antibiosis experiment (i.e., no-choice test) under laboratory conditions (26± 2°C, 60± 10% RH, and 12h photophase)

| Genotypes | Number of eggs/cladode<sup>a</sup> | Incubation period (hours)<sup>a</sup> | Hatchability (%)<sup>a</sup> | 1<sup>st</sup> instar duration (days)<sup>a</sup> | 2<sup>nd</sup> instar duration (days)<sup>a</sup> | Nymphal period (1<sup>st</sup> instar-Young female)<sup>a</sup> | 2<sup>nd</sup> instar weight (mg)<sup>a</sup> |
| --- | --- | --- | --- | --- | --- | --- | --- |
| Aakria | 130.0±25.4 a | 22.2±1.5 a | 82.8±11.9 a | - | - | - | - |
| Cherratia | 128.7±25.9 a | 22.3±1.7 a | 84.4±13.0 a | - | - | - | - |
| Marjana | 129.9±25.0 a | 22.3±1.6 a | 83.0±11.1 a | 37.0±2.2 a | - | - | - |
| Melk Zhar | 131.9±24.7 a | 21.7±1.8 a | 82.9±10.6 a | 36.1±2.1 ab | - | - | - |
| Angad | 132.8±24.0 a | 21.8±1.8 a | 83.2±10.0 a | 34.6±1.6 bc | - | - | - |
| B176 | 133.6±23.3 a | 22.2±2.0 a | 83.4±9.4 a | 30.1±1.8 cd | 43.5±1.1 a | 76.6±2.1 a | 4.2±0.4 b |
| B180 | 134.5±23.0 a | 21.4±1.5 a | 84.8±9.1 a | 32.7±1.9 cd | 42.9±1.5 ab | 75.6±2.3 ab | 4.3±0.5 b |
| Karama | 136.9±20.4 a | 21.6±1.3 a | 84.7±9.8 a | 32.3±1.5 d | 42.0±2.2 ab | 74.3±2.8 ab | 4.4±0.5 b |
| Ghalia | 137.8±19.8 a | 21.2±1.4 a | 84.1±9.6 a | 31.9±2.5 d | 41.6±2.3 b | 73.5±3.9 b | 4.4±0.5 b |
| Belara | 142.8±19.4 a | 20.9±1.5 a | 83.4±11.2 a | 27.6±3.2 e | 37.6±3.1 c | 65.1±3.9 c | 5.5±0.6 a |
| Statistics | $F = 0.69$<br>$df = 9, P = 0.71$ | $F = 1.91$<br>$df = 9, P = 0.05$ | $F = 0.10$<br>$df = 9, P = 1.00$ | $F = 36.29$<br>$df = 7, P < 0.0001$ | $F = 23.64$<br>$df = 4, P < 0.0001$ | $F = 44.08$<br>$df = 4, P < 0.0001$ | $F = 24.41$<br>$df = 4, P < 0.0001$ |

Means within a column followed by the same letters are not significantly different according to
Tukey's LSD test at $\alpha = 0.05$

For 1st instars, the genotypes Marjana (37 days), and Melk Zhar (36.1 days) induced the longest development times, whereas Belara induced the shortest development time for this stage. First instar nymphs fed Marjana, Melk Zhar, and A200 dying without having completed the 2nd nymphal stage. The B176 (43.5 days), B180 (42.9 days), and Karama (42 days) genotypes extended the development time of 2^nd^ instars, compared to Belara (37.6 days) genotype. The total nymphal development time (1st instar to young female) was significantly longer when *D. opuntiae* fed on B176 (76.6 days), B180 (75.6 days), and Karama (74.3 days) genotypes and shortest when nymphs fed on Belara (65.1 days) (Table 4).

Weights of 2^nd^ instars were significantly higher when the scale pests fed on Belara (5.5 mg) genotype compared to the other genotypes.

We recorded wide variation among genotypes in first instar nymph viability, which ranged from 0.0 % to 24.4 % (Fig 1). Higher rates of survival to second instar nymph stage were observed when first instar nymphs were fed Belara plants (24.4%). On the other hand, the genotypes Aakria (0.0%) and Cherratia (0.0%) negatively affected this parameter of the mealybug.

**Fig 1.**
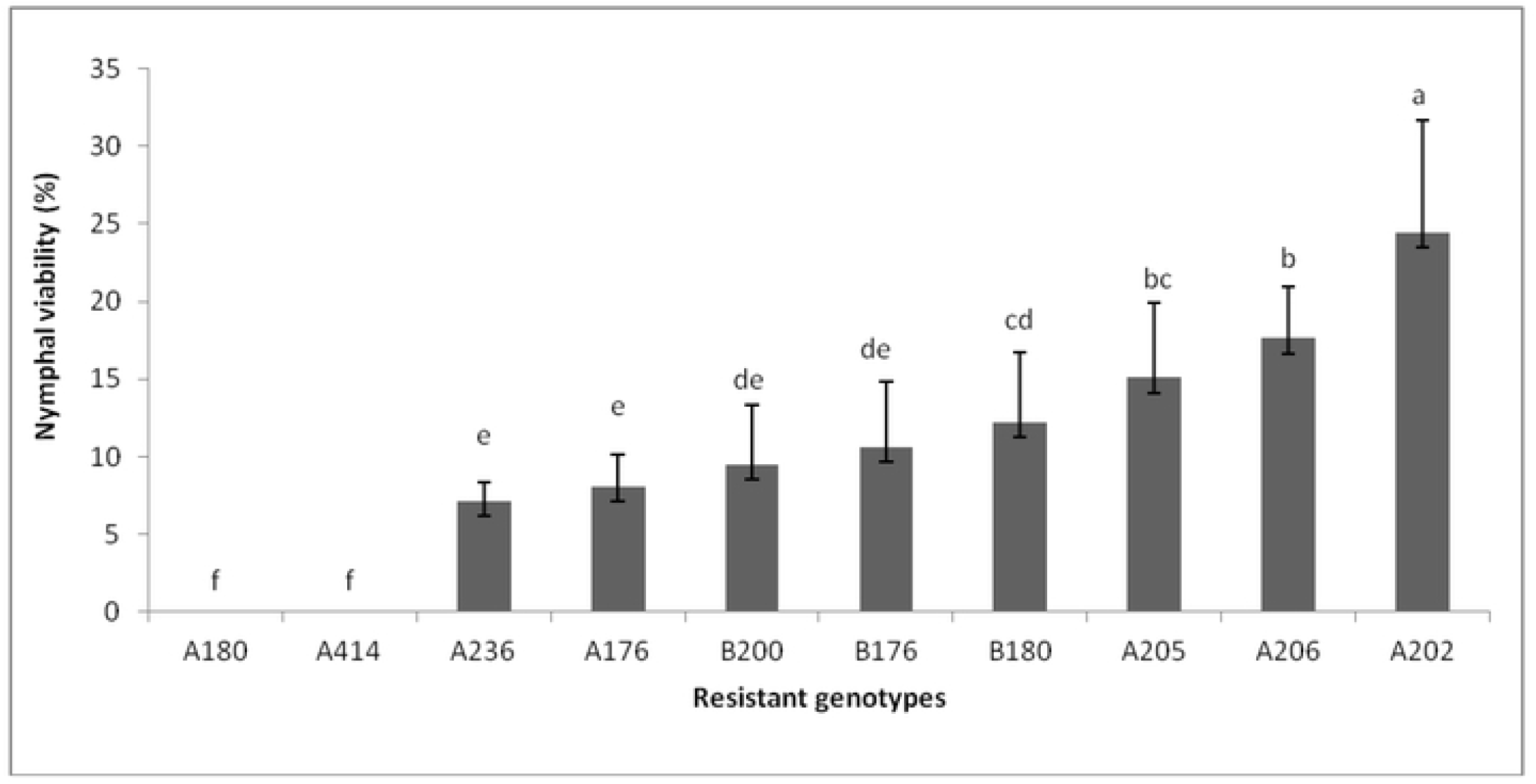
Nymphal viability (i.e., successful development to second instar nymph) of *D. opuntiae* on cactus resistant genotypes, in an antibiosis experiment (i.e., no-choice test). Different letters indicate significant difference between treatments by the Tukey’s LSD test at α= 0.05

#### Under greenhouse

The results are similar in shape to those obtained in the laboratory conditions, therefore confirming them. The results have shown that under greenhouse also the scale pest does not reach to develop definitively on the genotypes Aakria and Cherratia and doesn’t get beyond the young female stage in all the other tested resistant genotypes. Also, first instar nymphs fed Marjana, Melk Zhar, and A200 dying without having completed the 2^nd^ nymphal stage (Table 5). For the other genotypes tested, the nymphal period ranged from 54.7 to 65.5 days on average, with the longest mean development time on the genotypes B176, B180, and Karama and the shortest on Belara genotype (Table 5).

**Table 5.** Mean (±SE) development time of *Dactylopius opuntiae* reached stage on cactus genotypes in an antibiosis experiment under semi-field conditions (i.e., free-choice test)

| Genotypes | 1<sup>st</sup> instar duration (days)<sup>a</sup> | 2<sup>nd</sup> instar duration (days)<sup>a</sup> | Nymphal period<br>(1<sup>st</sup> instar- Young female)<sup>a</sup> |
| --- | --- | --- | --- |
| Aakria | - | - | - |
| Cherratia | - | - | - |
| Marjana | 32.1±2.2 a | - | - |
| Melk Zhar | 30.6±1.8 ab | - | - |
| Angad | 28.9±1.9 bc | - | - |
| B176 | 27.8±2.0 cd | 37.8±2.5 a | 65.5±3.5 a |
| B180 | 27.3±2.0 cd | 37.8±1.8 a | 65.1±2.3 a |
| Karama | 27.2±1.6 cd | 37.0±2.3 a | 64.2±2.6 a |
| Ghalia | 26.7±2.6 d | 36.4±2.5 a | 63.1±4.0 a |
| Belara | 22.2±3.6 e | 32.5±3.0 b | 54.7±4.1 b |
| Statistics | $F = 33.16$<br>df = 7, $P < 0.0001$ | $F = 16.03$<br>df = 4, $P < 0.0001$ | $F = 35.11$<br>df = 4, $P < 0.0001$ |

Means within a column followed by the same letters are not significantly different according to Tukey's LSD test at $\alpha = 0.05$

#### Morphological characterization of cactus genotypes

Principal component analysis of morphological traits of spines, cladodes, and whole plants showed clear groupings among cactus genotypes that correspond to their phylogenetic relationships. Karama and Ghalia shared cladode thickness and number of areoles with Belara. (Cherratia, Angad), and (Melk Zhar, Marjana), and (Aakria, B180) appeared as a distinct grouping respectively separated by the four morphometric characters tested (Fig 2).

**Fig 2.**
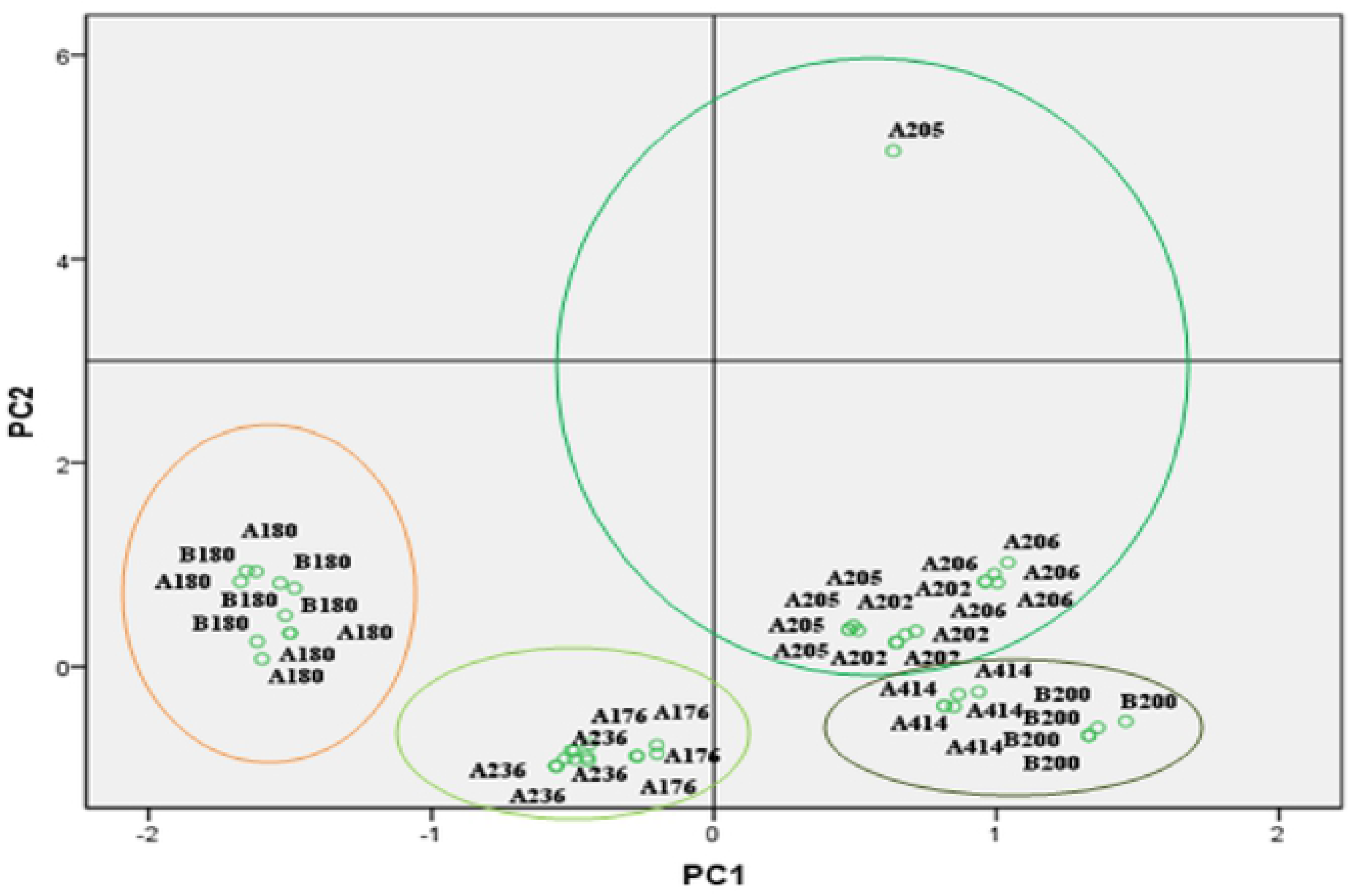
A principal component analysis (PCA) using four morphometric characters (i.e. total number of spines per cladode, cladode surface area, number of areoles per cladode, and number of areoles per plant) for the ten cactus genotypes identified as resistant to *D*.*opuntiae* in Morocco (n = 5 plant/genotype). A236: Marjana; A202: Belara ; A205: Karama ; A206: Ghalia; B200: Angad; A180: Aakria; A176: Melk Zhar; A414: Cherratia

## Discussion

Among the newly introduced enemies on cactus in Morocco, there is the mealybug “*Dactylopius opuntiae*”; a very devastating insect that is spreading in the Mediterranean countries with catastrophic damage on the cactus crop.

The management of this pest in the world, and more particularly in Morocco, is based mainly on several components, namely, i) the use of treatments by insecticides that have shown their limits in the field, ii) uprooting and burying of infested plants which is heavy, expensive and sometimes difficult to apply at the level of cactus plantations in the rough terrain or as hedges around houses, and finally, iii) scientific research. This last section focused on knowledge of this new pest, investigation of alternative methods of control, and research for genetic sources of cacti resistant to the wild scale “*Dactylopius opuntiae*”. Indeed, the first two axes have allowed enrichment of scientific knowledge useful for the continuation of the work related to the control of this new pest, while the third axis on the varietal resistance of cactus to the mealybug, was the relevant and saving solution. Thus, initially, the ten genotypes identified resistant by research in Morocco, will constitute a solid foundation for the launch of the national program of recovery of cactus decimated across the country by the Ministry of Agriculture, Maritime Fishing, Rural Development, and Water and Forests [23]. Our study documents the first study on the life history of *D*.*opuntiae* on the ten genotypes identified as resistant in Morocco using antexenosis and antibiosis tests in the laboratory and greenhouse.

Generally, antixenotic resistance to insects occurs due to the existence of morphological and/or chemical factors [46]. The results of this study (under laboratory and greenhouse conditions) indicated that all resistant genotypes tested showed a different level of antixenosis compared to the susceptible control and Aakria and Cherratia showed a strong antexenosis effect toward *D. opuntiae* (0-0.3 *D. opuntiae* alive 30 after infestation). The mechanisms underlying antixenosis resistance in these genotypes remain unknown. However, volatile chemical compounds may have differed between genotypes, which could determine whether a genotype will be more or less infested [47]. In addition, plants have the ability to produce multiple insecticidal compounds to defend themselves [41]. A new work published by Matos et al. (2021) that compared the chemical profile of four species of forage palms in Brazil (only one of which is susceptible to *D*.*opuntiae* and the others resistant), reported a total of 28 metabolites of which 18 were annotated [30]. The same authors indicated that quercetin, kaempferol, and isorhamnetin derivatives are distinguished as the main components of forage palm. Quercetin rhamnosyl dihexoside, quercitrin-3-O-2’,6’- dirhamnosylglucoside, and isorhamnetin-3-sophoroside 7-rhamnoside are the biomarkers that may be associated with resistance to *D. opuntiae* [30]. However, to confirm this hypothesis, additional studies should be performed.

No influence of cactus genotypes on insect biological parameters, including number of eggs per cladode, incubation period, and hatching percentage, was observed, indicating that the scale pest does not prefer any genotype over the others for oviposition. The mealybug fails to develop successfully on genotypes Aakria and Cherratia and does not develop beyond the young female stage in all other resistant genotypes tested. Also, first instar nymphs fed Marjana, Melk Zhar and A200 died without reaching the second instar nymphal stage, in addition, all the genotypes tested prolonged nymphal development of *D. opuntiae* and adversely affected nymphal viability (<24%), indicating resistance (antibiosis and/or antixenosis) to the cactus mealybug.

Antixenosis and antibiosis typically overlap, meaning that genotypes with high levels of antixenotic determinants may also have deleterious effects on insect life history, causing similar effects to plants that express antibiosis [31, 32]. For this reason, it may be difficult to differentiate between the two categories of resistance [48, 49] and may require specific determination of the mechanisms involved in insect toxification and detoxification. To avoid this misconception, new tools have been developed. For stink bugs, for example, electrical penetration graph (EPG) techniques have recently begun to be used to characterize feeding behavior [50, 51] and may be used in the future to characterize plant resistance categories. Second instar weights were significantly higher on the Belara genotype (5.5 mg) and no differences were observed among the other genotypes. In the present study, we observed that Belara leads to faster development of the nymphal stages of *D. opuntiae* compared to the other genotypes in which the insect successfully develops to the young female stage.

## Conclusion

The results of this study showed that the genotypes Aakria and Cherratia were the least attractive to *D*.*opuntiae*, indicating the expression of a strong antixenosis effect towards the scale pest. Under multiple choice conditions, the mealybug preferred no one genotype over the others for oviposition. Under laboratory and semi-field conditions, mealybug failed to develop on the genotypes Aakria and Cherratia and did not grow beyond the young female stage on all other resistant genotypes tested. In addition, the first instar nymphs fed on genotypes Marjana, Melk Zhar, and A200 died without reaching the second instar nymphal stage. In addition, all genotypes tested prolonged nymphal development of *D. opuntiae* and negatively affected their viability (<24%), indicating resistance (antibiosis and/or antixenosis) to the cactus mealybug. Genotypes Aakria and Cherratia showed the greatest stability of resistance as they showed a high level of antibiosis and antixenosis effect. Considering the extensive damage caused by *D. opuntiae* to cactus crops worldwide, further studies are needed to better interpret the resistance factors shown by certain genotypes. This information will be useful for breeding programs focused on pest resistance, in addition to assisting in the management of *D. opuntiae* in cactus plantations.

